# Investigation of Molecular Weights and Pharmacokinetic Characteristics of Older and Modern Small Drugs

**DOI:** 10.1101/2022.09.21.508888

**Authors:** Urban Fagerholm

## Abstract

**Background:** A shift towards higher molecular weight (MW) of drug candidates is anticipated to lead to changed pharmacokinetics (PK), including deteriorated absorption.

**Methods:** The objective of the study was to investigate changes in MW and PK of drugs over time by comparing MW and measured PK of small drugs (here MW<1500 g/mole) marketed before 2010 (n=277) and MW and *in silico* predicted (data produced using the ANDROMEDA by Prosilico software) and of small drugs marketed in 2021 (n=28).

**Results:** Apparently, there has been a shift towards higher MW (from 355 to 551 g/mole on average). This has influenced PK-parameters such as unbound fraction (on average approximately halved), fraction excreted renally (on average approximately halved; markedly decreased contribution by active secretion), bile excretion (almost 4-fold increased appearance; now for more than every other drug) and intrinsic metabolic clearance (increased). The very high percentage of modern drugs with (according to *in silico* predictions) significant renal and biliary excretion and gut-wall extraction, metabolic stability, limited passive intestinal permeability+efflux, limited gastrointestinal dissolution/solubility potential and/or a very low f_u_ increases complexity in predictions and places demands on predictive laboratory and computational methods.

**Conclusion:** Increased MW and changed PK-profiles (increased complexity) with time were observed. This shows the need for updating method set-ups for quantification and prediction of PK-parameters. ANDROMEDA has the capability to predict and optimize human clinical PK-characteristics of modern drug candidates with high accuracy.

## Introduction

The qualitative concept *druglikeness* used in drug design describes how “druglike” compounds are with respect to different factors, such as the pharmacokinetic (PK) factors fraction absorbed (f_a_) and oral bioavailability (F). Various methods for *PK-druglikeness* have been proposed and used, including Lipinski’s *Rule of 5*. According to this rule, an orally bioavailable drug candidate or drug has no more that one violation of the following 4 criteria: ≤5 hydrogen bond donors, ≤10 hydrogen bond acceptors, a molecular weight (MW)≤500 g/mole, and a log P (a lipophilicity measure) not exceeding 5 (1). The *Rule of 5* was extended to the *Rule of 3* for defining compounds with lead-like properties, and for this rule, narrower criteria were set: ≤3 hydrogen bond donors, ≤3 hydrogen bond acceptors MW≤300 g/mole and log P≤3 (2).

During drug discovery, log P and MW are often increased in order to improve the affinity and selectivity of drug candidates, and a result may be poorer and/or changed PK-characteristics. Consequences of increases of MW are (generally) decreased permeability, solubility and gastrointestinal uptake and increased renal and biliary clearance (CL_R_ and CL_bile_) and impact of efflux (3–6), which in turn makes the PK more complex and difficult to predict.

The objective of the study was to investigate changes in MW and PK of drugs over time by comparing MW and measured PK-profiles of small drugs (here defined as MW<1500 g/mole) marketed before 2010 and MW and predicted (data produced using the ANDROMEDA by Prosilico software) and observed PK-profiles of drugs marketed in 2021.

## Methods

### Data sets and parameters

The human PK data bank for pharmaceutical drugs marketed up to year 2010 compiled by Varma et al. (7) was selected as the older reference data set. PK parameters in this set included unbound fraction in plasma (f_u_), total clearance (CL), hepatic CL (CL_H_) and CL_R_. Out of 309 compounds 277 had reported fu-values, and these 277 were selected for this study. Fraction excreted via the renal route (f_e_) was calculated as CL_R_/CL, CL_R_/(GFR•f_u_)≥1.5 defined contribution by active renal secretion, CL_R_/(GFR•f_u_)≤2/3 defined contribution by tubular reabsorption, and intrinsic CL (CL_int_) was calculated using CL_H_, f_u_ and the well-stirred model with a hepatic liver blood flow rate of 1500 mL/min. Additional bile excretion data were taken from various literature sources.

Human PK-data for 28 small drugs marketed in 2021 (5 of which are dosed intravenously and have more complete PK-profiles) were mainly selected from FDA (Food and Drug Administration)-label documents (Clinical Pharmacology sections) of each drug (see also 8). The amount of human PK-data for the new small drugs was, however, limited, with data for only 1 to 6 compounds for many parameters (8).

### *In silico* predictions of PK

*In silico* predictions of human clinical PK of drugs marketed in 2021 were produced using ANDROMEDA by Prosilico software. Descriptions of the software and results can be viewed in reference 8.

## Results

### Observed data

Median MWs for older and modern small drugs are 334 and 477 g/mole, respectively (Figure 1). Corresponding mean MWs are 355 and 551 g/mole, respectively. 7 and 46 % of older and modern drugs have a MW exceeding 500 g/mole, respectively.

**Figure 1.**
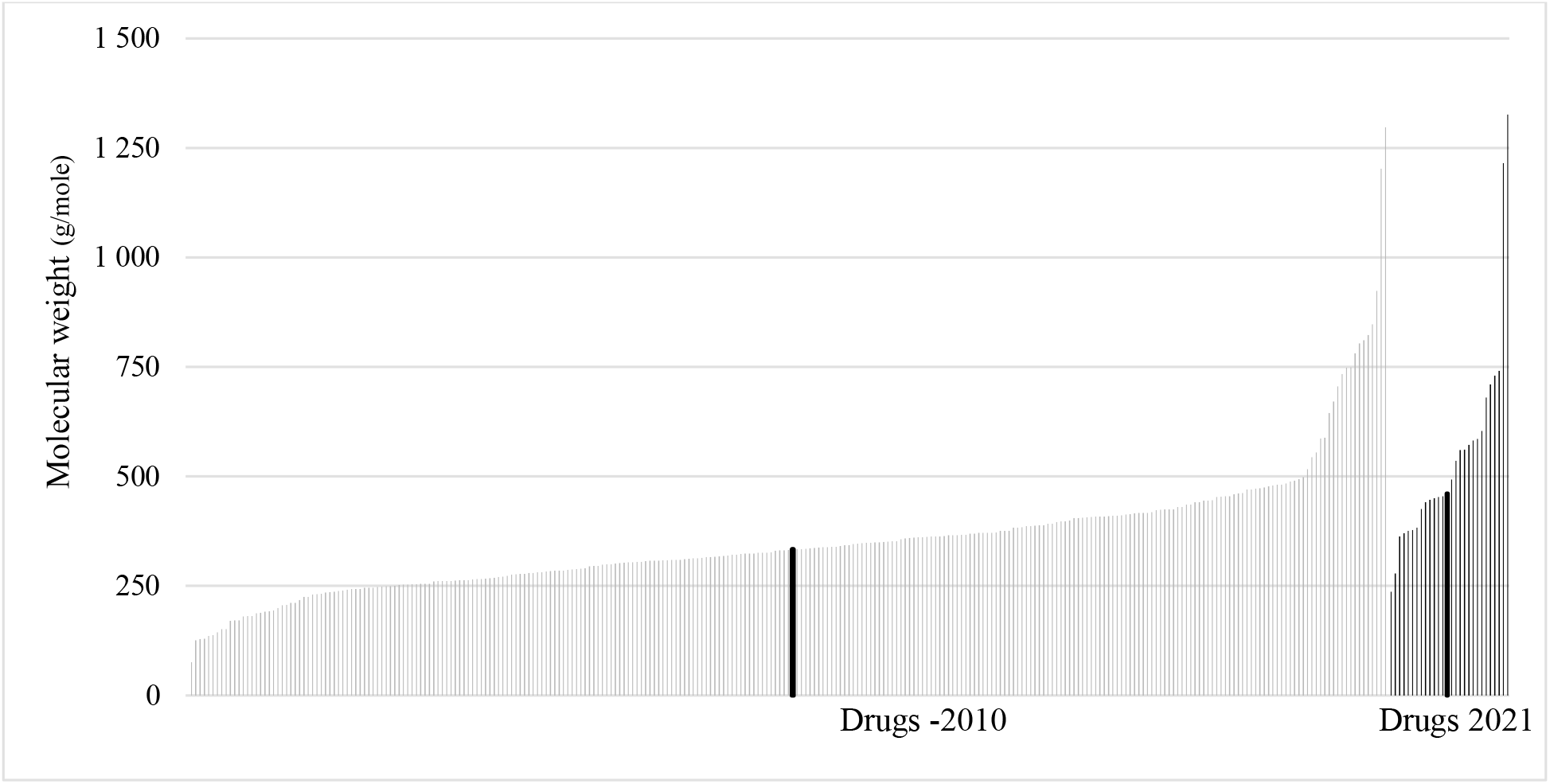
Molecular weights of small drugs marketed up to 2010 and during 2021. The thick bars note median values (334 g/mol for small drugs marketed up to 2010 and 477 g/mole for small drugs marketed in 2021).

Median and mean f_u_ and f_e_ for older drugs are 0.24 and 0.37, and 0.17 and 0.33, respectively (Figure 2). Corresponding f_u_- and f_e_-estimates for modern drugs are lower, 0.04 and 0.19, and 0.11 and 0.18, respectively. 19 % of older drugs and 41 % of modern drugs have a f_u_≤0.02, 48 % of older drugs and 25 % (3 of 12 with available data) of modern drugs have a f_e_≥0.2, and 32 % of older drugs and 8 % (1 of 12 with available data) of modern drugs have a f_e_≥0.5.

**Figure 2.**
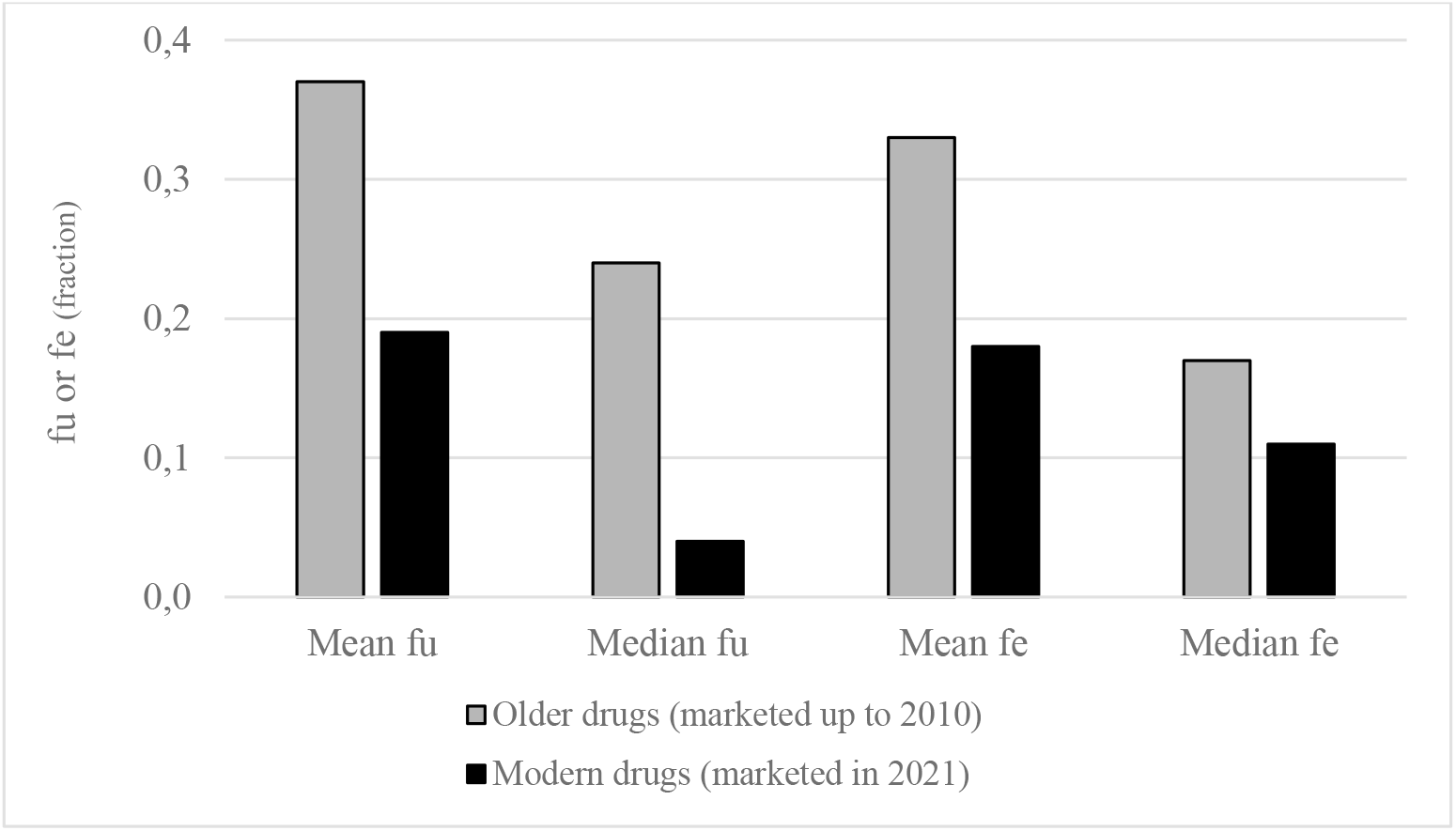
Mean and median unbound fraction in plasma (f_u_) and fraction excreted renally (f_e_) for older and modern drugs.

CL_R_ for 42 and 36 % of older drugs involves active secretion and reabsorption, respectively. Only 3 modern drugs have available CL_R_-data, and 2 of these (67 %) seem to be actively secreted and none to have apparent contribution by reabsorption.

Median and mean CL_int_ for older drugs are 676 and 22010 mL/min, respectively (Figure 2). For the modern small drugs only 2 CL_int_-values are available (24 and 8750 mL/min). 54 % of CL_int_-values for older drugs were below the corresponding limit of quantification (LOQ) of the conventional human hepatocyte assay (<1000 mL/min).

12 % of older drugs were identified as being excreted in bile. For drugs launched in 2021 there are at least 2 (7%) compounds suspected for bile excretion.

### Predicted data

According to *in silico* predictions, 29, 46, 54, 46, 43, 43, 32, 29 and 4 % of small drugs marketed in 2021 are significantly eliminated via the renal route (f_e_≥20%) and in the gut-wall (F_gut_<80%), are excreted via bile, have contribution by tubular reabsorption, have low/moderate passive intestinal permeability+efflux, have metabolic stability (CL_int_<1000 mL/min; corresponding to LOQ of the human hepatocyte assay), have limited gastrointestinal dissolution/solubility potential, have very low f_u_ (≤0.02) and are actively secreted in the kidneys, respectively (Figure 3) (8).

**Figure 3.**
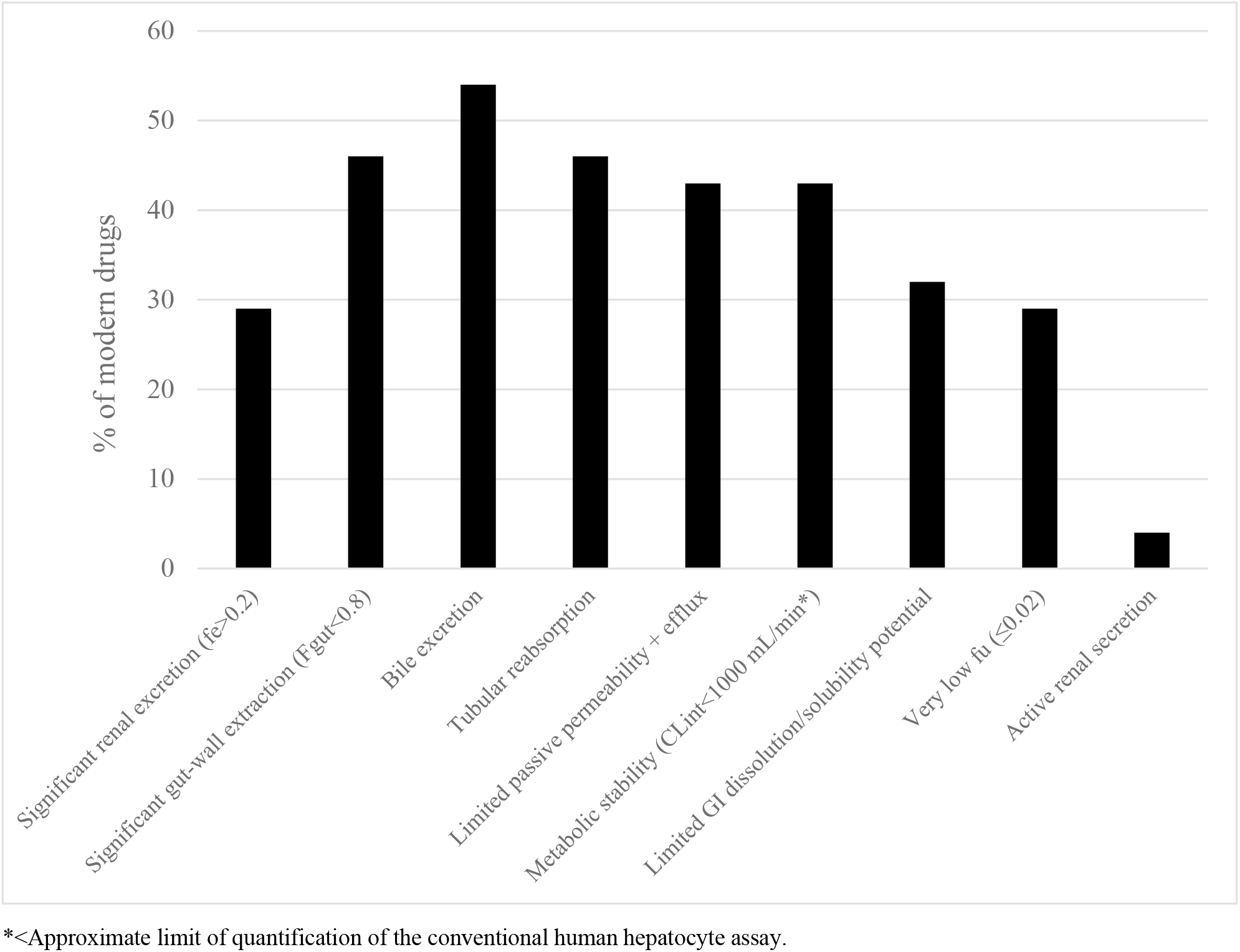
Percentages of new small drugs marketed in 2021 with certain *in silico* predicted PK-characteristics (8).

No compound had a combination of all these. 25 and 11 % of them had a combination of renal and bile excretion and metabolic stability or renal and bile excretion, metabolic stability, gut-wall extraction and limited gastrointestinal dissolution/solubility potential, respectively. According to predictions, only 1 of 28 small modern drugs had no significant absorption limitation, gut-wall extraction and excretion.

Predicted median and mean CL_int_ for modern drugs are 1402 and 7893 mL/min, respectively. 43 % of CL_int_-values were below the corresponding LOQ of the conventional human hepatocyte assay (<1000 mL/min).

## Discussion

Apparently, there has been a shift towards higher MW for small drugs since before 2010 (average from 355 to 551 g/mole; +55 % on average; now 46 % of drugs with MW>500 g/mole), and this has influenced PK-parameters such as f_u_ (on average approximately halved), f_e_ (on average approximately halved; cases of active secretion reduced to 1/10) and bile excretion (almost 4-fold increased appearance; now for more than every other drug) and CL_int_ (roughly doubled median CL_int_ and lower fraction below 1000 mL/min, but reduced mean value, according to observed data for older drugs and *in silico* predictions for modern drugs). The increased presence of excretion in bile is consistent with the trend towards increased MW (6).

The very high percentage of modern drugs with significant renal and biliary excretion, gut-wall extraction, metabolic stability, limited passive intestinal permeability+efflux, limited gastrointestinal dissolution/solubility potential and/or a very low f_u_ is a challenge for laboratories producing data for human clinical PK-predictions (9–11). For example, at low f_u_ there is increased uncertainty/variability for f_u_ and PK-predictions (9), and this is also the case for compounds with low Pe and S (10).

Despite having MWs above 500 g/mole the modern compounds have reached the market as medicines for oral administration. This indicates that *Rule of 5* may too strict for defining non-druglikeness. This has been highlighted and discussed earlier, by for example Hagan et al (12), who found that only about every other orally administered new chemical entity obeys it.

An advantage with ANDROMEDA by Prosilico is its capability to predict the human clinical PK for all the new small drugs with high accuracy, including parameters such as f_u_, f_e_, CL_R_, CL_bile_, F_gut_ and *in vivo* dissolution potential, fa and F for compounds with low S and Pe, and CL_int_ for relatively stable compounds. Median, mean and maximum prediction errors for a set-up of various PK-parameters were 2.5-, 3.5- and 16-fold, respectively (8). The prediction accuracy with ANDROMEDA by Prosilico was on par with (12 % of comparisons) or better than (76 % of comparisons) with the best laboratory-based prediction methods and the prediction range was considerably broader.

## Conclusion

Increased MW and changed PK-profiles (increased complexity) with time were observed. This shows the need for updating method set-ups for quantification and prediction of PK-parameters. ANDROMEDA has the capability to predict and optimize PK-characteristics of modern drug candidates with high accuracy.

## References

1. Lipinski CA, Lombardo F, Dominy BW, Feeney PJ. Experimental and computational approaches to estimate solubility and permeability in drug discovery and development settings. Adv Drug Del Rev. 2001;6:3–26.

2. Congreve M, Carr R, Murray C, Jhoti H. A ‘rule of three’ for fragment-based lead discovery? Drug Discov Today. 2003;8:876–877.

3. Waring MJ. Defining optimum lipophilicity and molecular weight ranges for drug candidates - Molecular weight dependent lower log D limits based on permeability. Bioorg Med Chem Letters 2009;19:2844–2851.

4. Fagerholm U. Prediction of human pharmacokinetics - Renal metabolic and excretion clearance. J Pharm Pharmacol. 2007;59:1463–1471.

5. Fagerholm U. The role of permeability in drug ADME/PK, interactions and toxicity - Presentation of a permeability-based classification system (PCS) for prediction of ADME/PK in humans. Pharm Res. 2008;25:625–638.

6. Fagerholm U. Prediction of human pharmacokinetics - Biliary and intestinal clearance and enterohepatic circulation. J Pharm Pharmacol. 2008;60:535–542.

7. Varma MVS, Obach RS, Rotter C, Miller HR, Chang G, Steyn SJ, El-Kattan A, Troutman MD. Physicochemical space for optimum oral bioavailability: Contribution of human intestinal absorption and first-pass elimination. J Med Chem. 2010;53:1098–1108.

8. Fagerholm U, Hellberg S, Alvarsson J. Spjuth O. *In silico* predictions of human clinical pharmacokinetics with ANDROMEDA by Prosilico – Predictions for a proposed benchmarking data set and new small drugs on the market 2021 and comparison with laboratory methods. Accepted for publication in ATLA.

9. Fagerholm U, Spjuth O, Hellberg S. Comparison between lab variability and *in silico* prediction errors for the unbound fraction of drugs in human plasma. Xenobiot. 2021;51:1095–1100.

10. Pham-The H, Garrigues T, Bermejo M, González-Álvarez I, Cruz Monteagudo M, Ángel Cabrera-Pérez M. 2013. Provisional classification and *in silico* study of biopharmaceutical system based on Caco-2 cell permeability and dose number. Mol Pharmaceut. 10:2445–2461.

11. Fagerholm U, Spjuth O, Hellberg S. The impact of reference data selection for the prediction accuracy of intrinsic hepatic metabolic clearance. J Pharm Sci. 2022;111:2645–2649.

12. Hagan SO, Swainston N, Handl J, Kell DB. A ‘rule of 0.5’ for the metabolite-likeness of approved pharmaceutical drugs. Metabolomics 2015;11:323–339.

